# Stochastic simulation platform for visualization and estimation of transcriptional kinetics

**DOI:** 10.1101/825869

**Authors:** Gennady Gorin, Mengyu Wang, Ido Golding, Heng Xu

## Abstract

We present an implementation of the Gillespie algorithm that simulates the stochastic kinetics of nascent and mature RNA. Our model includes two-state gene regulation, RNA synthesis initiation and stepwise elongation, release to the cytoplasm, and stepwise degradation, a granular description currently tractable only by simulation. To facilitate comparison with experimental data, the algorithm predicts fluorescent probe signals measurable by single-cell RNA imaging. We approach the inverse problem of estimating underlying parameters in a five-dimensional parameter space and suggest optimization heuristics that successfully recover known reaction rates from simulated gene expression turn-on data. The simulation framework includes a graphical user interface, available as a MATLAB app at https://data.caltech.edu/records/1287.

## Introduction

Transcription has been the focus of intensive study due to its cornerstone role in cell activity regulation. Recent advances in fluorescent imaging have enabled mRNA detection at single-molecule resolution in individual cells, in both live and fixed samples (1,2). Spatial analysis of mRNA signals allows the identification (3,4) and quantification (5) of nascent (actively transcribed) mRNA, which offers a direct window into the kinetics of gene transcription, with minimal interference from downstream effects (5), at the level of a single gene copy (6).

Converting high-resolution experimental data into theoretical understanding of transcription requires simultaneous modeling of both nascent and mature species of mRNA. Particularly, since at any given moment an mRNA molecule may be in a partially transcribed and/or degraded state, a good model should be able to capture the submolecular features of mRNA. However, current computational models of transcription present challenges for integration with the new wealth of microscopy data. Most models do not distinguish between nascent and mature mRNA or model the transcript length (7–11). As recently noted (5), several mechanistic models do describe the elongation of nascent mRNA, but do not consider the mature mRNA population and require additional processing for comparison to microscopy data (4,12–14). Further, studies using these models tend to predict low-order statistics (7,13), which paint a limited picture at biologically low molecule numbers (4,15). Recent methods based on directly solving the chemical master equation (CME) (5,15,16), using the finite state projection (FSP) algorithm [13], yield distributions of the number of molecules. However, integrating the discrete CME with submolecular features of mRNA is nontrivial, and has only recently been accomplished on a model with a deterministic elongation process (5). A stochastic stepwise model of transcription, more faithful to the mechanistic details, is not currently tractable using FSP (5) due to exponential growth in the size of the state space with increasing resolution.

Here we present a stochastic simulation platform that aims to capture the complexities of RNA processing. The platform consists of a submolecular implementation of the Gillespie algorithm (17), simulating the gene switching, transcription, and degradation expected in a prokaryotic system. Transcription and degradation occur in a stochastic fashion, where the initiation and individual steps of elongation are Poisson processes. The algorithm outputs time-dependent fluorescent probe signals, calculated from the overlap of intact RNA and probe-covered regions. The probe signals are provided as cell-specific readouts and as aggregated histograms, mimicking live-cell (MS2) and fixed-cell (smFISH) fluorescence data, respectively (1,2). Using a GUI, a user can input simulation parameters and examine time-dependent statistics, as well as animate the instantaneous molecule states.

We use the platform to approach the inverse problem of biological parameter estimation. A recent investigation demonstrated that entire distributions are required to reliably estimate parameter values from single-cell mRNA data (15). To perform parameter estimation based on these empirical distributions, we implement a heuristic approach based on iteratively minimizing mean squared errors and Wasserstein distances of different observables (18). This approach represents a novel method of estimating plausible regions for multiple parameters using time-series data with multiple observables, without making assumptions regarding the functional form of the distributions. Thus, the platform provides a flexible simulation environment to implement reaction mechanisms as well as a search algorithm designed to directly test those mechanisms’ parameters against experimental data. The GUI and search algorithm are available at https://data.caltech.edu/records/1287.

## Results

### Model and simulation platform

Our platform models a common formalism for the mRNA transcription process (5,7), with a series of stochastic reactions, including promoter turn-on and turn-off, transcription initiation, elongation, RNase (ribonuclease) binding, and degradation (**S1 Table**). Specifically, promoter activity is represented as a two-state switch. In the active (“on”) state, transcription can be initiated. The nascent mRNA strand elongates from the 5’ to the 3’ end, in a series of discrete steps. Upon reaching the end of the template gene, the mature mRNA molecule is released from the gene. Regardless of RNA maturity, RNase can bind to the 5’ end of the mRNA, causing the strand to begin stepwise degradation at an average rate assumed to be identical to the elongation speed (19). The process is depicted in **Fig 1A**. The physiology of the transcribed gene is parametrized by the turn-on rate *k*_*on*_, the turn-off rate *k*_*off*_, the transcription initiation rate *k*_*ini*_, the degradation initiation rate *k*_*deg*_, the elongation speed *v*_*el*_, and the gene length *L*. The experimental parameters include the timespan of the experiment *T*_*end*_, as well as the probe span vector (*P*_3_, *P*_5_) defining its 3’ and 5’ limits of coverage with respect to the length of the gene, as shown in **Fig 1A** (5).

**Fig 1.**
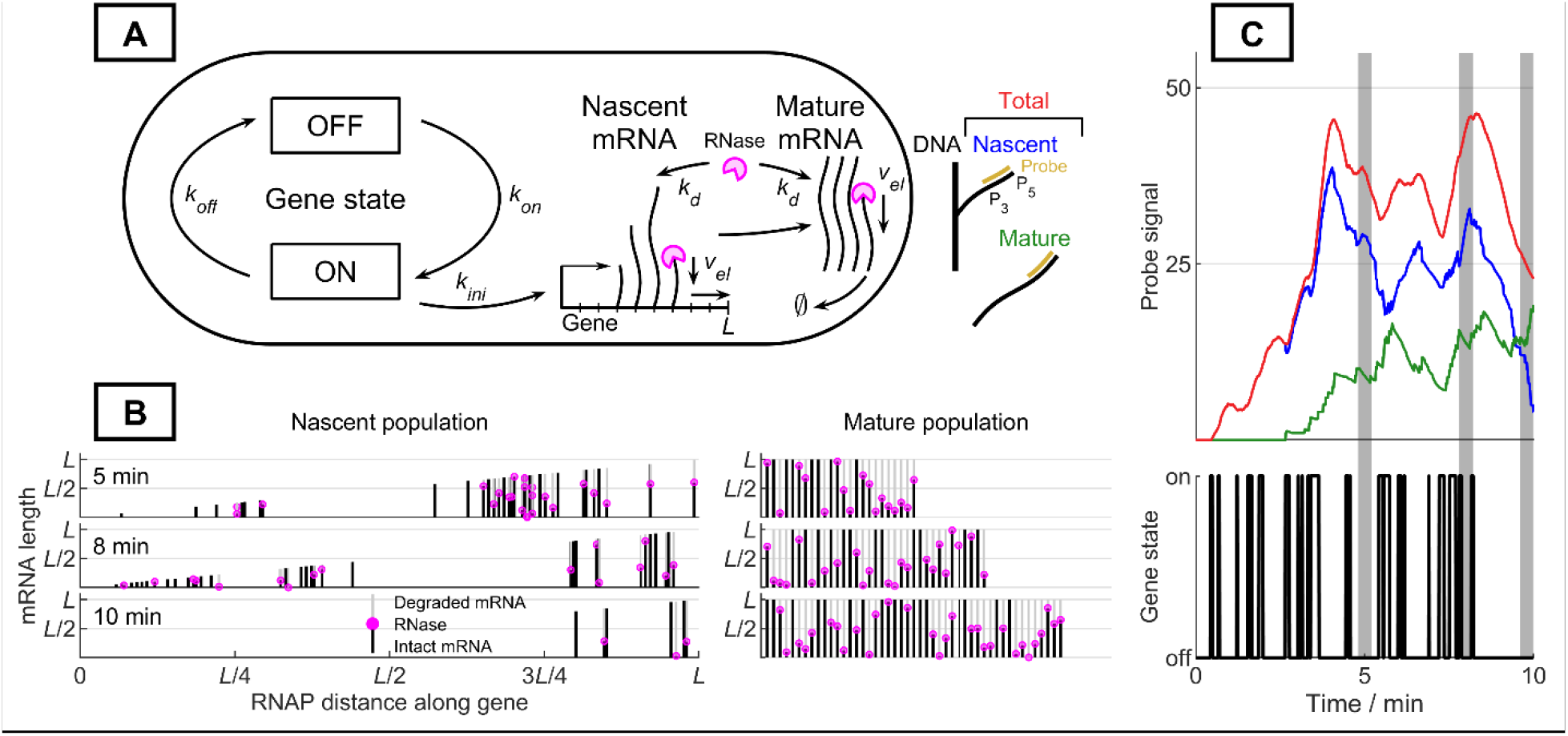
Model and simulation platform. **A:** Model schematic and probe parameterization (gold: probe coverage, *P*_3_: 3’-most edge of the probe, *P*_5_: 5’-most edge of the probe) **B:** Time-dependent molecule-level visualizations available through the GUI. Trajectory generated using *k*_*ini*_ = 100 min^−1^, *k*_*on*_ = 3 min^−1^, *k*_*off*_ = 10 min^−1^, *k*_*deg*_ = 0.5 min^−1^, *v*_*el*_ = 41.5 nt s^−1^, *T*_*end*_ = 10 min, *L* = 5300 nt, 241 steps of elongation to complete transcription (dark line: intact RNA stretches, light line: degraded RNA stretches, pink circle: RNase molecule). **C**: Single-cell trajectory with simulated nascent and mature fluorescent signals. Parameters same as in **B** (red: total signal, blue: nascent signal, green: mature signal, shaded regions: times displayed in **B**).

The platform performs stochastic simulation of the model using the Gillespie algorithm (17,20), then estimates the fluorescence of each mRNA molecule from the size of its region targeted by fluorescent probes. Specifically, we simulate the production and degradation of each mRNA molecule in the cell, whose status can be defined by four variables, i.e. two integers that define 5’- and 3’-most nucleotides of the transcript and two Boolean variables that define whether the mRNA is polymerase-bound (nascent) and/or RNase-bound (degrading). The gene state (on or off) is defined by a single Boolean variable. To convert the simulated mRNA molecule ensemble (**Fig 1B**) to the experimentally observed fluorescent signal, we calculate the overlap between the intact RNA and the probe coverage (single realization shown in **Fig 1C**); the probe readout is rescaled to molecule number using the fluorescence of a single intact molecule (16). The resolution of the simulation is determined by the number of cells and the number of steps taken to fully elongate or degrade each molecule.

Model simulation is implemented in MATLAB 2018a (21). A simple graphical user interface (GUI), provided as a MATLAB app at https://data.caltech.edu/records/1287, runs the simulation for a user-defined parameter set defining the physical parameters and simulation precision. Upon completion, the GUI outputs the time-dependent mean probe signal (in units of molecule number), Fano factor, and instantaneous nascent and total mRNA probe signal histograms, all calculated over the cell population. The mRNA nucleotide spans are used to visualize and animate the transcriptional activity taking place at an individual gene copy (analogous to **Fig 1B** and **Fig 1C**; example visualization given in **S1 Movie**). Our software allows direct simulation of complex experimental designs. For instance, to mimic the commonly-used induction experiment (e.g. the addition of isopropyl β-D-1-thiogalactopyranoside, an inducer of the *lac* promoter, to *E. coli* cells (6)), the simulation starts with no mRNA and undergoes a step increase in the gene turn-on rate. Similarly, to mimic a repression experiment (e.g. the addition of 2-nitrophenyl-β-D-fucoside to *E. coli*), the system starts with a steady-state population of mRNA and undergoes a step decrease in gene turn-on rate (22). For physiologically plausible transcription in short, infrequent bursts (23), the decrease in *k*_*on*_ can also model repression by a step decrease in initiation (6) caused by the addition of rifampicin (24).

### Parameter estimation

Given single-cell time-series fluorescence data that describes nascent and mature mRNA, we seek to estimate the underlying model parameters. We would like to approach this inverse problem by simulating mRNA number distributions for the experimentally available timepoints, evaluating an error metric that maps the divergence between the target distribution and each trial distribution to a single number, then minimizing this error by using it as an objective function.

Since metrics based on noisy empirical stochastic distributions do not meet the smoothness assumptions of gradient-based optimizations methods (25), we select a genetic algorithm for optimization. We use the MATLAB implementation of the genetic algorithm (21,26) to sample and evolve points in a parameter space spanning several orders of magnitude for each variable. Consistent with previous investigations, we use a logarithmic parameter search space (15). Each trial parameter vector {*k*_*ini*_, *k*_*on*_, *k*_*off*_, *k*_*deg*_, *v*_*el*_} is evaluated using an ensemble of hundreds to thousands of simulated cells. Due to the high computational load (millions of cell trajectories) of a single search, we vectorize the computation and parallelize it across processors on the Amazon Web Services (AWS) cloud (27). Since cells are independent, the algorithm scales well by parallelization across multiple processors. At the end of the simulation, the parallelized cell ensemble is reassembled into a single population and the statistics defining the error are computed locally, as shown in **Fig 2A**. To speed up convergence to consistent parameter sets, our heuristic method uses a variable objective function, with five distinct stages that use different error metrics. Details of the metrics are provided in **Methods**.

**Fig 2.**
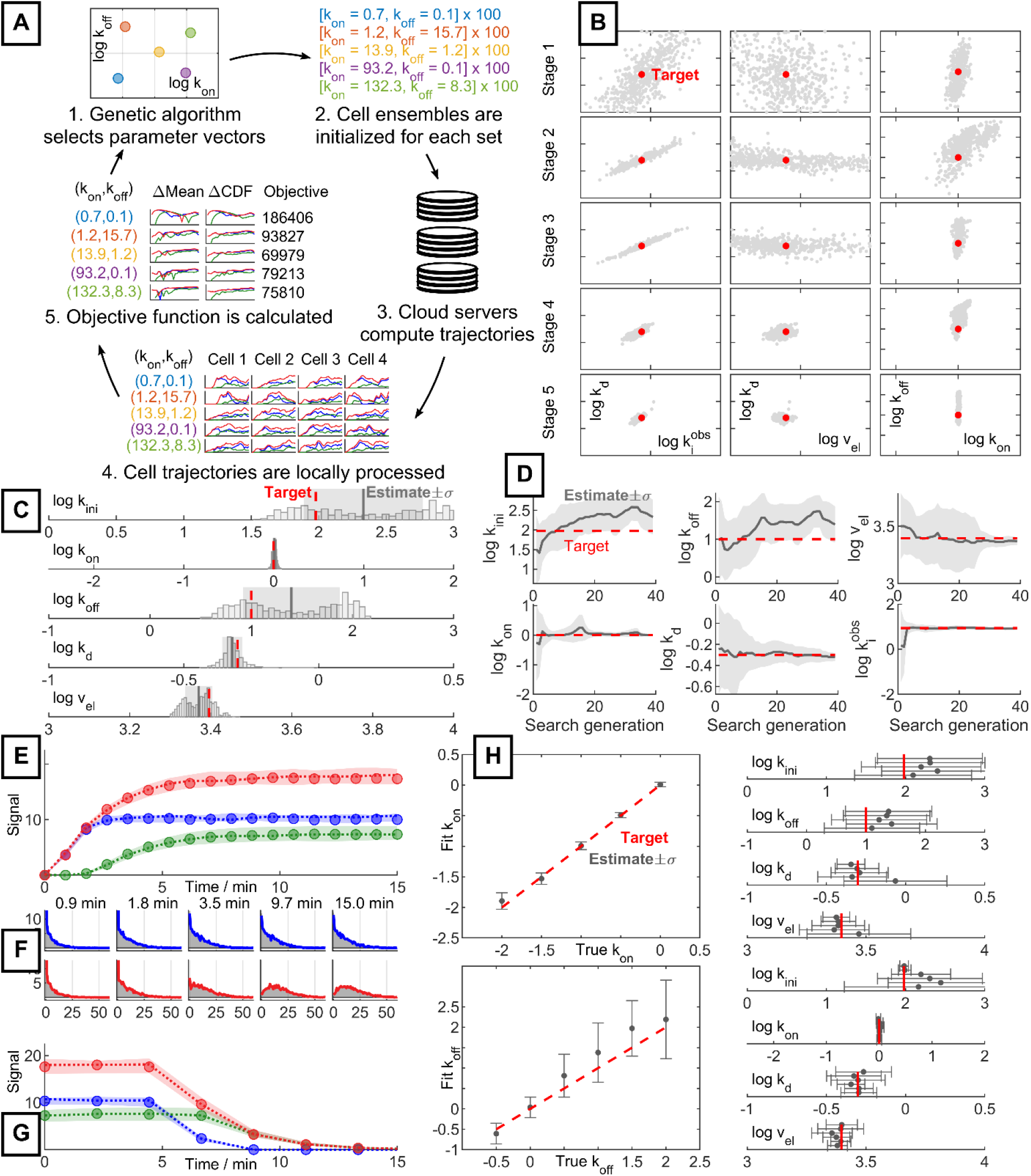
Parameter estimation process and performance. **A:** Parallelized calculation of the search objective function for a set of trial parameters (ΔMean: mean squared error, ΔCDF: Wasserstein distance, Objective: error function value). **B:** Convergence of the genetic algorithm (red: ground truth target, gray: population of parameter estimates). **C:** Final trial parameter population from **B** (red: ground truth target, histogram: estimate population, gray line: mean estimate, gray region: one-sigma region of estimates). **D:** Evolution of parameter estimates throughout the search process (red: ground truth target, gray line: mean estimate, gray region: one-sigma region of estimates). **E:** Comparison of mean probe signal between target and fit (circles: target data, dotted line: mean parameter estimate, shaded region around dotted line: signal spanned by fifty estimates sampled from the one-sigma region). Colors as in Fig 1. **F:** Comparison of copy-number distributions between target and fit (shaded gray regions: target histogram, colored lines: histogram generated from mean parameter estimate, top row/blue: nascent mRNA distribution, bottom row/red: total mRNA distribution). **G:** Comparison of mean probe signal between target and fit in turn-off cross-validation experiment. Convention as given for **E**. **H:** Estimation of modulated parameters. Top trial modulates k_on_, bottom trial modulates k_off_. All other parameters are constant but unknown to the search algorithm and are fit independently (red: ground truth target, gray dots and error bars: mean estimate and one-sigma region of three replicates).

To test the algorithm’s ability to recover known parameters, we generated synthetic data for the turn-on experiment using the following ground truth parameters: *k*_*ini*_ = 95 min^−1^, *k*_*on*_ = 1 min^−1^, *k*_*off*_ = 10 min^−1^, *k*_*deg*_ = 0.5 min^−1^, *v*_*el*_ = 41.5 nt s^−1^, *T*_*end*_ = 15 min, *L* = 5300 nt, 10,000 cells, and 15 steps of elongation to complete transcription. The procedure used to convert these rates into reaction propensities is described in the **Supplementary Information**. Relatively coarse simulation quality was used as a proof of concept.

The simulations were parallelized across 90 AWS processors. The process of parameter identification is visualized in **Fig 2B**. We found that the one-sigma interval around the mean estimate included the ground truth parameters (**Fig 2C**). The convergence of *k*_*on*_, *k*_*deg*_, and *v*_*el*_ throughout the search is relatively well-behaved and close to monotonic; however, *k*_*off*_ and *k*_*ini*_ are far more challenging to estimate (**Fig 2D**).

We compare the mean signals of nascent and total RNA simulated using the one-sigma estimate interval (**Fig 2E**), as well as the corresponding distributions simulated using the mean estimate (**Fig 2F**), to the synthetic ground truth data. Comparison at both levels demonstrates convergence. To cross-validate the search, we compare repression simulations generated from the ground truth and estimated parameters. The nascent and total means are consistent (**Fig 2G**). To test the robustness of the fitting algorithm, we apply the search procedure to the turn-on data generated using a range of *k*_*on*_ and *k*_*off*_ values, mimicking the regulatory parameter modulation hypothesized to occur *in vivo* (28). The results suggest consistent performance throughout the parameter space, although identifiability of high *k*_*off*_ is poor (**Fig 2H**). Encouragingly, all one-sigma intervals include the ground truth parameters.

## Methods

The Gillespie algorithm is adapted from the original description (17) and implemented in the MATLAB programming language (21). To account for submolecular degrees of freedom, the simulation uses multiple data structures to describe the system state. Specifically, one multidimensional dynamic array holds the 5’ and 3’ indices of each mRNA (transcript span), another identifies whether it is being transcribed at a particular gene locus or free in the cytoplasm (RNA polymerase attachment), and a third tracks whether it is being degraded (RNase attachment). Smaller, static arrays track the system time, gene state, and number of mRNA and bound RNase molecules. Each reaction either increments or flips Boolean values in the appropriate state arrays. State variables and reactions are outlined in detail in **Supplementary Information**; the reaction propensity calculations are given in **S1 Table**.

To perform parameter estimation on turn-on synthetic data, we use a heuristic iterative method based on the genetic algorithm (29). We alternate between optimizing mean signals and entire distributions. The error metric for the mean signal is the mean squared error. Due to the limited support of empirical distributions, the commonplace minimization of Kullback–Leibler divergence between target and test distributions (30) is inappropriate for comparing distributions (25). Instead, we use the absolute difference between the target and test cumulative distribution functions (CDFs), which tends to be more robust to noise and sparsity (25); this metric is commonly known as the Wasserstein or earth mover’s distance (18). We aggregate different time points’ Wasserstein distances by weighing them using a uniform or exponential function of time, as described in **Supplementary Information**.

Empirically, the parameter identifiability is far from uniform throughout the simulated time-series, and different metrics provide sensitivity to different parameters. Further, it is computationally prohibitive to simulate entire trajectories at the beginning of the parameter search, when the relevant region of the five-dimensional search space is not yet known. Therefore, we take an *ad hoc* iterative approach, which incrementally narrows the region of parameters consistent with the observed signals. This heuristic approach is chosen for computational convenience and is not guaranteed to the global parameter optimum.

The first stage identifies the parameter space consistent with the distributions of nascent signals observed throughout the first few time points of the experiment, essentially acting as an order-of-magnitude filter and eliminating computationally expensive edge regions with extremely high or low transcription. This stage uses a population of 5,000 parameter sets and only keeps the top 10% best variants; based on **Figure 2B**, it identifies *k*_*on*_ and a degenerate line containing consistent values of *k*_*d*_ and 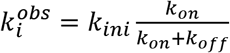. The second stage attempts to truncate this space to parameter values consistent with the mean level of total RNA for the entire time series, and identifies tighter bounds for *k*_*d*_ and *k*_*i*_^*obs*^. This and all following stages use populations of 500 trial parameters. The third stage refines the estimate to parameter values consistent with the steady state distribution of total mRNA, and yields tighter bounds for *k*_*on*_ and *k*_*off*_. The fourth stage uses information from the mean level of nascent RNA for the entire time series, and improves bounds for *v*_*el*_. Finally, the fifth stage refines the bounds for *k*_*on*_ and *k*_*off*_ by performing a high-precision optimization using the metric used in stage 1. By penalizing the objective function for deviating beyond a given radius from the previous stage’s parameter region, consistency between different error metrics is enforced, as described in **Supplementary Information**. More detailed data regarding each stage’s penalization and precision are provided in **S2 Table**.

## Discussion

Above we describe a new platform for simulating mRNA transcription and degradation on a submolecular level, available at https://data.caltech.edu/records/1287. Its output is directly comparable to single-cell data of nascent and mature mRNA. The output of each simulation is the empirical distribution of signals for each cell at each time point. Therefore, the platform can simulate both live-cell measurements (which identify cell-specific signals over time) and fixed-cell measurements (which yield population statistics) (1,2). As the platform is based on the stochastic simulation algorithm, it is relatively straightforward to modify the model to incorporate new reactions, chemical species, regulatory pathways, and labeling schema. The software includes single-cell and statistical visualization tools to facilitate general-purpose use without coding. For resource-intensive parameter space exploration, we suggest heuristics to accelerate convergence. The method demonstrates that parameter estimation from a time series of multiple observables is tractable by heuristic likelihood-free methods. The validation we perform suggests that, by using simulations to generate empirical distributions, this approach is more effective to fit experimental signals than traditional methods when no closed-form solutions or approximations are available; further, the visualization capabilities would be useful for the qualitative description and understanding of such complex systems.

Our platform allows numerical solution of detailed transcription model for both nascent and mature mRNA species, whose CME may not be solved exactly. However, since the approach is simulation-based, the steady state of the system needs to be computed asymptotically from a non-steady state, which may be time-consuming. Specifically, simulating and fitting the steady-state and turn-off experiments may be computationally prohibitive if the scales of kinetic rates have substantial difference. Alternatively, it may be possible to use analytical solutions (31,32) to approximate an equilibrium distribution; however, this approach is challenging to generalize and the resulting simulation would no longer be exact.

The parameter identification process may be facilitated by parameter constraints from analytical solutions. For example, if the steady-state solution for the total mean is known, the *k*_*i*_^*obs*^ and *k*_*deg*_ parameters can be fixed for the optimization procedure, reducing the parameter estimation to the simpler problem of optimization in three-dimensional space of *k*_*on*_, *k*_*off*_, and *v*_*el*_. However, we anticipate that the value of this heuristic method rests in applications to models with *ad hoc* mechanisms that do not have easily tractable analytical solutions.

The system is currently limited to modeling prokaryotic transcription. The implementation of eukaryotic transcription would require making significant changes to the reaction schema, such as disabling nuclear nascent degradation and adding a kinetic model of a transport process after the release of the newly transcribed mRNA. The implementation of more complex transcription schema, such as disallowing polymerase progression past another polymerase (33), which are challenging to theoretically treat using the CME framework, are relatively straightforward to achieve by checking the distance to the nearest neighbor in front of each polymerase and making the propensity of elongation zero if the distance is sufficiently small. The enforcement of delays at particular locations along the gene is likewise straightforward, specifically by replacing the constant elongation speed with a speed dependent on the polymerase location. Finally, time-dependent propensities, corresponding to phenomena such as cell cycle regulation, are trivial to implement, facilitating the extension of the platform to more complex systems.

## Supporting information

Transcription visualization

## Acknowledgments

G.G. is supported by the California Institute of Technology Division of Chemistry and Chemical Engineering and NIH U19MH114830. G.G. thanks Dr. Lior Pachter (California Institute of Technology) for support and Dr. Brian Munsky (Colorado State University) for valuable advice. M.W. and I.G. are supported by grants from the National Institutes of Health (grant no. R01 GM082837) and the National Science Foundation (grant nos PHY 1430124). H.X. is supported by the National Key R&D Program of China (grant no. 2018YFC0310800), the National Natural Science Foundation of China (grant no. 11774225), the Thousand Talents Plan of China (Program for Young Professionals), the National Science Foundation of Shanghai (grant no. 18ZR1419800), and the Burroughs Wellcome Fund Career Award at the Scientific Interface (grant no. 1013907). We gratefully acknowledge the computing resources provided by the student innovation center at Shanghai Jiao Tong University.

## Notes

https://data.caltech.edu/records/1287

## References

1. Golding I, Paulsson J, Zawilski SM, Cox EC. Real-Time Kinetics of Gene Activity in Individual Bacteria. Cell. 2005 Dec;123(6):1025–36.

2. Lee JH. Quantitative approaches for investigating the spatial context of gene expression: Spatial context of gene expression. WIREs Syst Biol Med. 2017;9(2):e1369.

3. Iyer S, Park BR, Kim M. Absolute quantitative measurement of transcriptional kinetic parameters in vivo. Nucleic Acids Res. 2016;44(18):e142–e142.

4. Zenklusen D, Larson DR, Singer RH. Single-RNA counting reveals alternative modes of gene expression in yeast. Nat Struct Mol Biol. 2008;15(12):1263–71.

5. Xu H, Skinner SO, Sokac AM, Golding I. Stochastic Kinetics of Nascent RNA. Phys Rev Lett. 2016;117(12):128101.

6. Wang M, Zhang J, Xu H, Golding I. Measuring transcription at a single gene copy reveals hidden drivers of bacterial individuality. Nat Microbiol [Internet]. 2019 Sep 16 [cited 2019 Sep 23]; Available from: http://www.nature.com/articles/s41564-019-0553-z

7. Munsky B, Neuert G, van Oudenaarden A. Using Gene Expression Noise to Understand Gene Regulation. Science. 2012;336(6078):183–7.

8. Paulsson J. Models of stochastic gene expression. Physics of Life Reviews. 2005 Jun;2(2):157–75.

9. Honkela A, Peltonen J, Topa H, Charapitsa I, Matarese F, Grote K, et al. Genome-wide modeling of transcription kinetics reveals patterns of RNA production delays. Proc Natl Acad Sci USA. 2015;112(42):13115–20.

10. Bokes P, King JR, Wood ATA, Loose M. Exact and approximate distributions of protein and mRNA levels in the low-copy regime of gene expression. J Math Biol. 2012 Apr;64(5):829–54.

11. Shahrezaei V, Swain PS. Analytical distributions for stochastic gene expression. Proceedings of the National Academy of Sciences. 2008 Nov 11;105(45):17256–61.

12. Kim S, Jacobs-Wagner C. Effects of mRNA Degradation and Site-Specific Transcriptional Pausing on Protein Expression Noise. Biophysical Journal. 2018;114(7):1718–29.

13. Choubey S. Nascent RNA kinetics: Transient and steady state behavior of models of transcription. Phys Rev E. 2018;97(2):022402.

14. Tripathi T, Chowdhury D. Interacting RNA polymerase motors on a DNA track: Effects of traffic congestion and intrinsic noise on RNA synthesis. Phys Rev E. 2008;77(1):011921.

15. Munsky B, Li G, Fox ZR, Shepherd DP, Neuert G. Distribution shapes govern the discovery of predictive models for gene regulation. Proc Natl Acad Sci USA. 2018;115(29):7533–8.

16. Skinner SO, Xu H, Nagarkar-Jaiswal S, Freire PR, Zwaka TP, Golding I. Single-cell analysis of transcription kinetics across the cell cycle. eLife. 2016 Jan 29;5:e12175.

17. Gillespie DT. A general method for numerically simulating the stochastic time evolution of coupled chemical reactions. Journal of Computational Physics. 1976 Dec;22(4):403–34.

18. Deza M, Deza E. Encyclopedia of distances. Dordrecht : New York: Springer Verlag; 2009. 590 p.

19. Chen H, Shiroguchi K, Ge H, Xie XS. Genome‐wide study of mRNA degradation and transcript elongation in Escherichia coli. Molecular Systems Biology [Internet]. 2015 [cited 2019 Sep 15];11(1). Available from: https://www.embopress.org/doi/abs/10.15252/msb.20145794

20. Gillespie DT. Exact stochastic simulation of coupled chemical reactions. J Phys Chem. 1977 Dec;81(25):2340–61.

21. MATLAB R2018a [Internet]. The MathWorks, Inc.; 2018. Available from: https://www.mathworks.com/products/matlab.html

22. Elf J, Li G-W, Xie XS. Probing Transcription Factor Dynamics at the Single-Molecule Level in a Living Cell. Science. 2007;316(5828):1191–4.

23. Dar RD, Razooky BS, Singh A, Trimeloni TV, McCollum JM, Cox CD, et al. Transcriptional burst frequency and burst size are equally modulated across the human genome. Proceedings of the National Academy of Sciences. 2012 Oct 23;109(43):17454–9.

24. Eron L, Block R. Mechanism of Initiation and Repression of In Vitro Transcription of the Lac Operon of Escherichia coli. Proc Nat Acad Sci USA. 1971;5.

25. Poovathingal SK, Gunawan R. Global parameter estimation methods for stochastic biochemical systems. BMC Bioinformatics. 2010;11(1):414.

26. MATLAB R2018a Global Optimization Toolbox [Internet]. The MathWorks, Inc.; 2018. Available from: https://www.mathworks.com/products/global-optimization.html

27. Amazon Web Services. AWS General Reference - Reference guide. 2019; Available from: https://docs.aws.amazon.com/general/latest/gr/aws-general.pdf

28. Sanchez A, Golding I. Genetic Determinants and Cellular Constraints in Noisy Gene Expression. Science. 2013 Dec 6;342(6163):1188–93.

29. Goldberg DE. Genetic Algorithms in Search, Optimization and Machine Learning. 1st ed. Boston, MA, USA: Addison-Wesley Longman Publishing Co., Inc.; 1989.

30. Munsky B, Fox Z, Neuert G. Integrating single-molecule experiments and discrete stochastic models to understand heterogeneous gene transcription dynamics. Methods. 2015;85:12–21.

31. Jahnke T, Huisinga W. Solving the chemical master equation for monomolecular reaction systems analytically. J Math Biol. 2006 Dec 11;54(1):1–26.

32. Schnoerr D, Sanguinetti G, Grima R. Approximation and inference methods for stochastic biochemical kinetics—a tutorial review. J Phys A: Math Theor. 2017 Mar 3;50(9):093001.

33. Kim S, Beltran B, Irnov I, Jacobs-Wagner C. Long-distance cooperative and antagonistic RNA polymerase dynamics via DNA supercoiling. Cell. 2019 Sep 19;179(1):106–119.

