## Supplementary Information for "Stochastic simulation platform for visualization and estimation of transcriptional kinetics"

#### Gillespie algorithm

The Gillespie algorithm is adapted from the original description (17,20). At a given simulation step, each cell has the choice of a characteristic set of $M$ reactions with corresponding propensities; an example set is provided below. The propensity of a reaction with the index $\mu \in \{1, \ldots, M\}$ is $a_\mu$, usually the mass action kinetic rate. The total propensity is $a$, defined as $\sum_{\mu=1}^{M} a_\mu$. The reaction index chosen is that $\mu$ which meets $\sum_{1}^{\mu-1} a_\nu < r_1 a \leq \sum_{1}^{\mu} a_\nu$, for a random number $r_1$ drawn from the uniform distribution on (0,1). Time is incremented by $\tau = (1/a) \ln (1/r_2)$, where $r_2$ is a different random number drawn from the same distribution.

If there are $Q$ different molecular species in the system, the standard version of the algorithm requires only an array of length $Q$ to keep track of the system state (17). However, the inclusion of partial molecule fragments necessitates more complex data structures. Their dimensions are defined by the following conventions:

- $N$: the number of cells being simulated, user-set integer.
- $Z$: the size of the mRNA allocation, dynamically increasing integer.
- $n\_genes$: number of gene copies simulated in the system, a user-set integer.
- $n\_chemical\_spaces$: number of molecular species considered distinguishable, a model-dependent integer. Practically, if a microscopy method can distinguish between the mature mRNA and nascent mRNA of all $n\_genes$ gene copies, $n\_chemical\_spaces = n\_genes + 1$.
- $L$: length of the gene in nucleotides, a model-dependent integer.

The state of the system is described by 8 variables:

1. $RNA\_indices$: $N \times Z \times 2$ integer array. This array defines the intact portion of each RNA molecule, as defined by the 3' and 5' index of each RNA. For a molecule in cell $n \in \{1, \ldots, N\}$, allocated index $z \in \{1, \ldots, Z\}$, the 3'-most nucleotide index $I_3 \in [0, L]$ is stored at the position $(n, z, 1)$ and the 5'-most nucleotide index $I_5 \in [0, I_3]$ is stored at the position $(n, z, 2)$. Transcription gradually increments the 3' index from nucleotide 0 to nucleotide $L$; degradation does the same to the 5' index. Not every entry of the array corresponds to an RNA in the system; some are empty (filled with a $NaN$ value)
2. $RNA\_identity$: $N \times Z \times n\_chemical\_spaces$ binary array. This array defines the identity or location of each RNA molecule. For a molecule in cell $n \in \{1, \ldots, N\}$, allocated index $z \in \{1, \ldots, Z\}$, and chemical space $S \in \{1, \ldots, n\_chemical\_spaces\}$, this array has a Boolean $TRUE$ value at the position $(n, z, S)$.
3. $RNA\_rnase\_binding$: $N \times Z \times n\_chemical\_spaces$ binary array. This array defines whether an RNase molecule is bound to a given mRNA. For an RNase attached to an mRNA in cell $n$, allocated index $z$, and chemical space $S$, this array has a Boolean $TRUE$ value at the position $(n, z, S)$.
4. $RNA\_fill$: $N \times Z$ binary array. This array specifies the mRNA allocation pattern. It is equal to the output of the MATLAB command $any(RNA\_identity, 3)$; if cell $n$ has a molecule allocated in index $z$, this array has a Boolean $TRUE$ value at $(n, z)$ .

5. $gene\_active\_state$: $N \times n\_genes$ binary array. This array specifies the Boolean state of the transcription sites, $TRUE$ for on and $FALSE$ for off. For gene indexed by integer $j \in [1, n\_genes]$ in the on state in cell $n$, this array has a Boolean $TRUE$ value at $(n, j)$.
6. $t$: $N \times 1$ float vector that holds the simulation time for each of the $N$ cells. The current simulation time for cell $n$ is stored at the position $(n)$.
7. $x$: $N \times n\_chemical\_spaces$ float vector that specifies the total number of molecules of each $n\_chemical\_spaces$ species, for all $N$ cells. If cell $n$ has $\xi$ molecules in chemical space $S$, this array has $\xi$ at the position $(n, S)$.
8. $rnase$: $N \times n\_chemical\_spaces$ float vector that specifies the total number of RNase molecules attached to molecules of each $n\_chemical\_spaces$ species, for all $N$ cells. If cell $n$ has $\eta$ RNase molecules attached to mRNA in chemical space $S$, this array has $\eta$ at the position $(n, S)$.

Due to space considerations, the first five variables are passed by reference, as fields of a custom class derived from the handle superclass. The last three are passed by value.

The model used here consists of eight reactions. The definitions use the following conventions.

- $k_{d,nas}$: nascent degradation initiation (RNase attachment) rate.
- $k_{d,mat}$: mature degradation initiation (RNase attachment) rate. Generally set equal to $k_{d,nas}$.
- $k_{elo}$: rate of a single step of elongation, calculated as $v_{el} \times N_{st} / L$.
    - $v_{el}$: elongation rate in nucleotides per unit time.
    - $N_{st}$: user-defined number of steps per gene.
    - $L$: gene length in nucleotides.
    - $Nuc$: $N_{st}/L$, the size of a single step.
- $k_{deg}$: rate of a single step of degradation. Generally set equal to $k_{elo}$.
- $n_{nas}$: number of nascent molecules at a gene.
- $n_{mat}$: number of mature molecules.
- $n_{nas,rnase}$: number of nascent molecules with RNase attached.
- $n_{mat,rnase}$: number of mature molecules with RNase attached.
- $G$: gene state. $TRUE$ if on, $FALSE$ if off.

All rates are in units of events per unit time. The reactions proceed as described in **S1 Table**.

| Reaction | Rate | Description |
|---|---|---|
| 1. Initiation | $G\ k_{init}$ | Allocate entry to new mRNA. Expand arrays if necessary. Increment nascent RNA by 1. |
| 2. Nascent elongation | $n_{nas}k_{elo}$ | Choose mRNA to elongate. Increment $I_3$ by $Nuc$. If $I_3 > L$, decrement nascent and increment mature count by 1. Reassign chemical space identifier to mature population. |
| 3. RNase attachment to nascent RNA | $(n_{nas} - n_{nas,rnase})k_{d,nas}$ | Choose mRNA to attach. Set RNase binding for molecule to $TRUE$. Increment nascent-bound RNase count by 1. |
| 4. Degradation of nascent RNA | $n_{nas,rnase}\ k_{deg}$ | Choose nascent mRNA to degrade. Increment $I_5$ by $Nuc$. If $I_5 > I_3$, reject reaction and do not increment time |
| 5. Gene turn-on | $(\neg G)k_{on}$ | Set gene state to $TRUE$. |
| 6. Gene turn-off | $G\ k_{off}$ | Set gene state to $FALSE$. |
| 7. RNase attachment to mature RNA | $(n_{mat} - n_{mat,rnase})k_{d,mat}$ | Choose mRNA to attach. Set RNase binding for molecule to $TRUE$. Increment mature-bound RNase count by 1. |
| 8. Degradation of mature RNA | $n_{mat,rnase}\ k_{deg}$ | Choose mature mRNA to degrade. Increment $I_5$ by $Nuc$. If $I_5 > L$, decrement mature RNA by 1 and reset all allocated space associated with molecule; set $I_3$ and $I_5$ to $NaN$. |

**S1 Table: Implemented stochastic reactions.**

The mRNA nucleotide index data is stored in the $RNA\_indices$ and must be combined with $RNA\_identity$ data and probe properties to generate probe fluorescence outputs. The outputs are only calculated at the Gillespie time steps immediately before reporting times. The following conventions are used:

- $P_5$: 5’ end of probe, user-defined.
- $P_3$: 3’ end of probe, user-defined.
- $idvec$: $n\_chemical\_spaces\ \times 1$ identity vector for a particular transcript. For the transcript in cell $n\ \in\ \{1, \ldots, N\}$ and at allocated index $z\ \in\ \{1, \ldots, Z\}$, this is identical to $RNA\_identity(n, z, :)$. Since each mRNA is in one and only one chemical space (an mRNA cannot be transcribed from two genes; once it matures, it is no longer nascent), $idvec$ is a Boolean vector with one TRUE value at position $S\ \in\ \{1, \ldots, n\_chemical\_spaces\}$, corresponding to its identity.
- $probe\_out$: $n\_chemical\_spaces\ \times 1$ float vector. Stores the fluorescence outputs. Initialized as a zero vector.

For a given transcript, the calculation proceeds as follows:

1. Calculate the 5' index of the probe-labeled region: $L_5 = max(P_5, T_5)$.
2. Calculate the 3' index of the probe-labeled region: $L_3 = min(P_3, T_3)$.
3. Check that the transcript region and the probe region overlap: $ovlpChk = (T_5 < P_3)\&(T_3 > P_5)$
4. Compute the probe contribution for the transcript: $signal = ovlpChk \times (L3 - L5)$.
5. Add the transcript's signal to the appropriate chemical space accumulator: $probe\_out = probe\_out + idvec \times signal$.

After all mRNA molecules present in the system are added, the signal is divided by $(P_3 - P_5)$ to rescale the measurement in the units of the number of transcripts. If a set of chemical spaces cannot be disambiguated by observation (e.g., two gene copies on the same DNA strand), the resulting observables are calculated by adding together the underlying chemical spaces' signals.

### Parameter search algorithm

The search procedure attempts to minimize a measure of statistical distance between posited test data and experimental target data. The definitions use the following conventions:

- $num\_t\_pts$: number of time points collected in the experiment.
- $n\_bins$: number of bins used to generate a histogram from the fluorescence observations at each step. By default, bins are centered on integers from 0 to 100.
- $n\_obs$: number of observables collected, such as two sets of data corresponding to nascent and total fluorescence.
- $n\_conds$: number of conditions or parameter sets tested.

The target data is stored in a $num\_t\_pts \times n\_bins \times n\_obs$ array. For a given time and observable, an entry in the second dimension gives the fraction of observations that fall into the particular bin. The test data is stored in a $n\_conds \times num\_t\_pts \times n\_bins \times n\_obs$ array. For each test histogram, we use the Wasserstein metric to calculate the distance from the corresponding target histogram, yielding an $n\_conds \times num\_t\_pts \times n\_obs$ array. The time points for a particular parameter set and observable are aggregated through a user-defined weighing function, giving a $n\_conds \times n\_obs$ array. Only one observable is selected at each stage of the search; this choice yields $n\_conds \times 1$ array that maps each parameter set to one error value. Since specific cell identities are not tracked, this target data definition corresponds to copy-number distributions from fixed (smFISH) microscopy observations.

We use the built-in MATLAB function $ga$ to execute the genetic algorithm. To force it to report degenerate results, $ga\ EliteCount$ is set to 0. The most intensive computations tend to occur at domain edges; to avoid their generation, the crossover function is set to $@crossoverheuristic$ and new individuals are only generated by crossover, never mutation. The crossover ratio varies with stage.

To ensure that the search does not "forget" the location of the solution subspace found in the previous stage, parameter combinations not in the vicinity of the population at the end of the previous stage are considered infeasible. To enforce this condition, we add a compensating factor to the error function. For example, one of the test parameter sets in stage 2 is the vector $p$, while the population surviving at the end of stage 1 is the set of vectors $S = \{S_i\}$. The raw error function value is $E(p)$ and the compensating factor is $C(p, S)$. The algorithm optimizes over the logarithm of the parameter space, generally over $1 - 4$ orders of magnitude in each parameter, so the Euclidian distance between $p$ and each of the vectors in $S$ is a small non-negative float. There exists a value $r$, the smallest such distance: $r = \min_i \|S_i - p\|$. If $r$ is smaller than a user-defined radius R, the compensating factor is $C(p, S) = 0$. If $r > R$, we make the compensating factor a rapidly growing function of $r$, here $C(p, S) = \exp[3(r - R)] - 1$. Alternatively, a simple step function (e.g. $C(p, S) = 10 \; \forall \; r > R$) can be used. The total error used in the optimization is $E_{tot}(p) = E(p) + C(p, S)$.

The settings used in each stage of the developed parameter estimation method, introduced in **Table 1**, are elaborated upon in **S2 Table**. The first stage does not adhere to a fixed number of cells, instead gradually increasing the number of cells and decreasing the number of trial parameter sets. Specifically, the first generation of the first stage is simulated using 2500 parameter sets and 150 cells, a relatively coarse simulation quality. The second selects the 625 best-performing parameter sets (as measured by the stage's error metric, Wasserstein distances of nascent RNA distributions, with time-dependent weighing by an exponential function), recombines them, and simulates them using 600 cells. The consequent stages use 900, 1200, and 1500 cells respectively, and iteratively trim the total number of parameter sets to 584, 542, and 500. All stages afterward use 500 parameter sets and do not modulate the number of sets.

| Stage | Target | Metric | Time | Weighing | Gen. | Cells | Radius | Crossover ratio |
|---|---|---|---|---|---|---|---|---|
| 1 | Nas. | CDF | 0 to 3.5 min | $\exp(-10t/T_{end})$ | 5 | 1500 | N/A | 0.5 |
| 2 | Tot. | Mean | 0 to 15 min | Uniform | 8 | 500 | 0.14 | 0.25 |
| 3 | Tot. | CDF | 0 to 15 min | $\exp(3[t/T_{end} - 1])$ | 8 | 700 | 0.12 | 0.23 |
| 4 | Nas. | Mean | 0 to 15 min | Uniform | 8 | 900 | 0.11 | 0.16 |
| 5 | Nas. | CDF | 0 to 3.5 min | $\exp(-10t/T_{end})$ | 5 | 3500 | 0.12 | 0.16 |

**S2 Table. Parameter estimation settings, detailed.** Nas.: Nascent mRNA. Tot.: Total mRNA. CDF: Wasserstein metric of empirical cumulative distribution function. Mean: Total squared error of the mean. Weighing: Time-dependent weight multiplied by the metric to generate a scalar objective function value. Gen.: Number of generations of genetic algorithm. Cells: number of cells used in the simulation. Radius: R, the radius defining the plausible region about each candidate point.

### Source code

The MATLAB source code available with the article contains the following four files.

- *gillespie*172: Core simulation algorithm. Inputs parameters for physiology, simulation quality, and experiment type and timespan; outputs arrays corresponding to nascent, cytosolic, and total RNA probe signals, as well as an array of ground truth molecule numbers.
- *HandleState_int*: Helper function to implement pointers in MATLAB.
- *gillespie_target_generator_ex*1: Example function to generate a single ensemble time-series dataset.
- *gill_cell_ga_search_*1: Sample search algorithm to fit the dataset; outputs genetic algorithm population properties throughout the search.

### Graphical User Interface

The graphical user interface is implemented as the MATLAB app *RNA_growth_app*. The default view, after running a simulation, is shown in **Figure S1**.

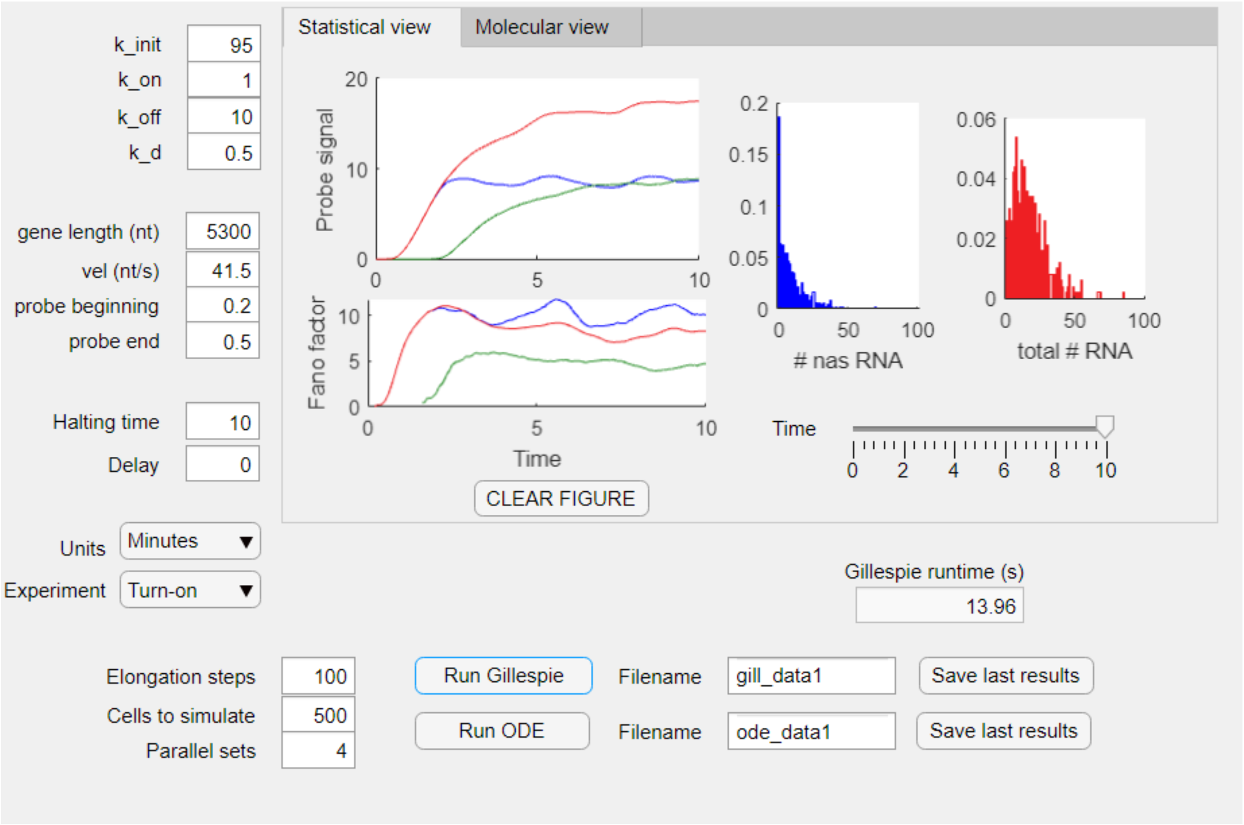


**Figure S1: MATLAB app, statistical view.**

The left side of the app defines the physical parameters and experimental setup. The top four fields correspond to rates of initiation, turn-on, turn-off, and degradation, in user-defined units. The next four fields correspond to gene and probe properties; probe beginning and end are numbers between 0 and 1 that define the span of the probe with respect to the transcript. Halting time is the length of the experiment. Turn-on, turn-off, and steady-state experiments are supported. Turn-on experiments start with no mRNA and experience a step increase in the turn-on rate after the delay. The other types start with equilibrated mRNA; turn-off experiments experience a step decrease in the turn-on rate after the delay.

The bottom-left fields define the simulation properties. If multiple cores are available on the computer and the MATLAB Parallel Computing Toolbox is installed, type in the number of cores to use into the "Parallel sets" field.

The buttons on the bottom allow simulation and data storage. Press "Run Gillespie" to execute the stochastic simulation. Press "Run ODE" to visualize the differential equation solution for comparison. The most recent simulation and ODE results are saved in memory; press the corresponding "Save last results" button to store them to disk under the desired filename. The field above displays the runtime for the most recent stochastic simulation.

The “Statistical view” panel permits the visualization of moments and distributions once data has been generated. The left two panels show the mean probe signal and Fano factor for nascent (blue), mature (green), and total (red) RNA. The right two panels depict empirical copy-number distributions for nascent and total RNA; the slider below is used to display the distribution at a desired timepoints. A “CLEAR FIGURE” button is available below the mean and Fano factor views. The button only erases these aggregated plots; distributions are erased automatically at each run. Simultaneous plotting is useful for qualitative visualization of the variation between successive runs, difference between signals for different probe spans, the effect of low vs. high cell numbers, the bias introduced by highly coarse simulations, and other comparative purposes.

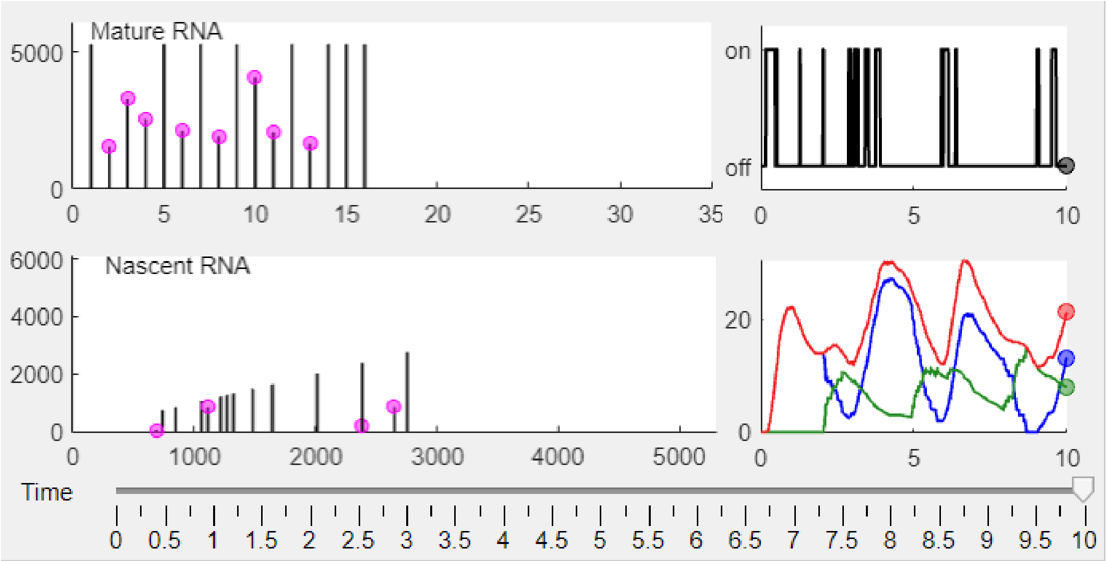


**Figure S2: MATLAB app, molecular view panel.**

The “Molecular view” panel (**Figure S2**) qualitatively visualizes growing and degrading mRNA in a single cell; all colors and conventions are as given in **Figure 1B** and **1C**. The top left panel shows the entire mature population. The abscissa is the integer index of each mRNA molecule; the ordinate is its remaining length. Purple dots represent RNase molecules and indicate that the mature mRNA is being degraded. The bottom left panel depicts the entire nascent population. The abscissa is distance along the DNA strand; the ordinate is the mRNA length. The top right panel shows the ground truth gene state trace as a function of time. The bottom right panel shows the probe signals observed from the cell as a function of time. The slider below the panels is used to display the available data for a desired timepoint.
